# The Phase Shift Correlation function uncovers periodic Ulcerative Colitis microbiome relationships without temporal sampling

**DOI:** 10.1101/2020.07.01.174409

**Authors:** Robert E. Brown, Alena Arlova, Patrick M. Gillevet

## Abstract

Correlation analysis is a fundamental technique to determine potential relationships within biological processes. However, many biological processes have been shown to function in a periodic manner. When modeling correlations, the fluctuations that are associated with periodicity cause significant issues. We have implemented a Phase Shift Correlation (PSC) algorithm, with a corresponding value PSCrho, to address the periodicity and phase variance associated with features that vary with the same frequency -- but are phase shifted. The phase shift could well indicate causality with one feature’s quantitative change leading to the change in the other feature.

We applied the PSC algorithm to Ulcerative Colitis (UC) microbiome data and compared the resulting feature relationships with the equivalent Spearman correlation function results. PSC located many instances of higher phase shifted correlations, where the corresponding Pearson correlation was low.

## Introduction

Many publications have addressed the periodic nature of biological networks, with the circadian rhythm as a major example [1,2]. One way to address the periodic nature of biological functions is to perform temporal studies where multiple samples are collected per participant. Several methods have been developed to discover and describe periodicity in systems and gene expression networks when time data or cell cycle phase is available [3-5].The downside of this approach is that without a priori knowing the frequency of the cycle one does not know the number of samples necessary to determine the periodicity of the function. Additionally, if the periodicity is short (< day) then the researcher must collect many samples in a day. However, time or cycle phase is not always available, specifically in datasets that have been collected for non-temporal studies. The technique to address feature (metabolites, microbiome, etc.) relationships in this manner has lagged.

### Linear correlation

algorithms, Pearson’s and Spearman (for non-parametric data), are key approaches in studying a correlation relationship between two features [6]. However, with the periodic nature of many biological functions, linear correlations may miss key relationships. Biological processes frequently will follow a path were the result of one Feature A rising or falling will impact abundance of Feature B, that will go on to impact Feature C. One artifact of using linear correlations appears as a perplexing scenario frequently seen in a correlation network diagrams – we call it the *correlation triangle paradox* (Fig 1). It occurs when Feature A is significantly correlated to Feature B, and Feature C is significantly correlated to Feature B, however, Feature A and feature C are not significantly correlated. Another disadvantage with linear correlations is that they cannot infer causality, as correlation does not indicate causation.

**Fig 1.**
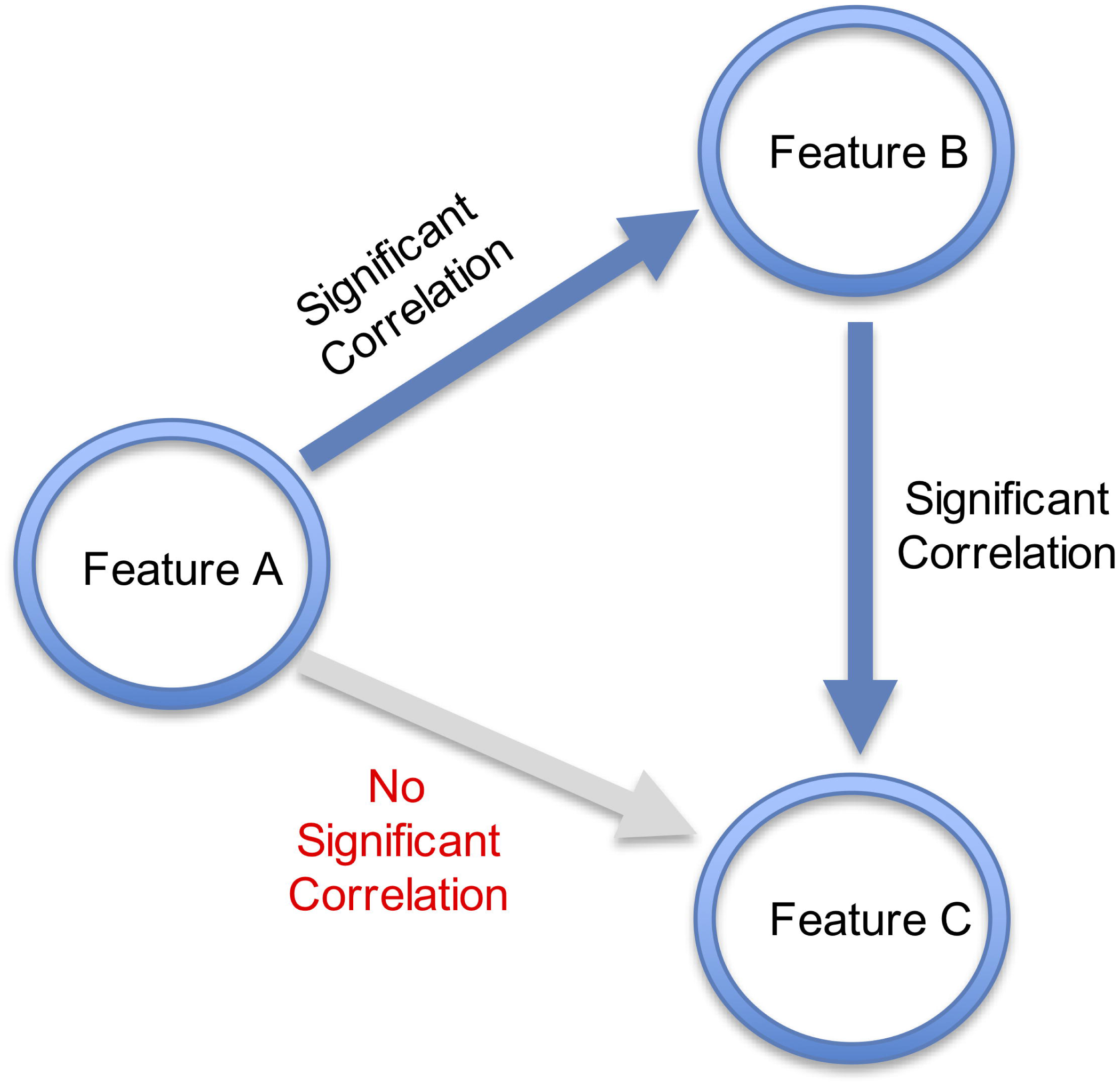
The correlation triangle paradox. Features A is significantly correlated to Feature B, and Feature B is significantly correlated to Feature C, however Feature A and Feature C are not significantly correlated.

We address this quandary using a Phase Shift Correlation (PSC) function on feature value snapshots based on a periodic function as opposed to applying a linear correlation function. The PSC identifies the same in-phase (zero-degree lag) positive and negative correlations as Pearson’s rho. However, if there is a phase shift (lag) between features, where the lag feature correlates with the lead feature by some phase difference, then the PSC may indicate the possibility of the lead feature having a potential causal relationship with the lag feature. Phase shift correlation affords researchers the opportunity to analyze complex dynamic biological systems as functions of time without requiring temporal data collection. As with linear correlations, phase shift correlations are an appropriate input for application of correlation difference probability networks for knowledge discovery by comparing significant changes in control correlation networks versus disease correlation networks [7].

## Materials and methods

The PSC approach does not directly address the actual periodicity of the phase delay between features but applies a cosine function between pairs of normalized feature values to calculate the applicability of an in-phase, phase shifted, or anti-phase relationship. The correlation values are derived from defining the error distance from the actual value of Feature A and the actual value of Feature B versus the estimated value for Feature B_est_ based on Feature A. The analysis is then repeated with Feature B as the lead feature. A key assumption in the PSCrho determination is assuming the possible feature specific maximum and minimum values across all samples are relatively constant. Then we can conclude that the value of each feature measured, here microbial abundance, in a study is an amplitude value along the vertical axis of a sinusoidal wave (Fig 2). Plus, changes to one feature (microbe) may directly impact the values of other feature(s).

**Fig 2.**
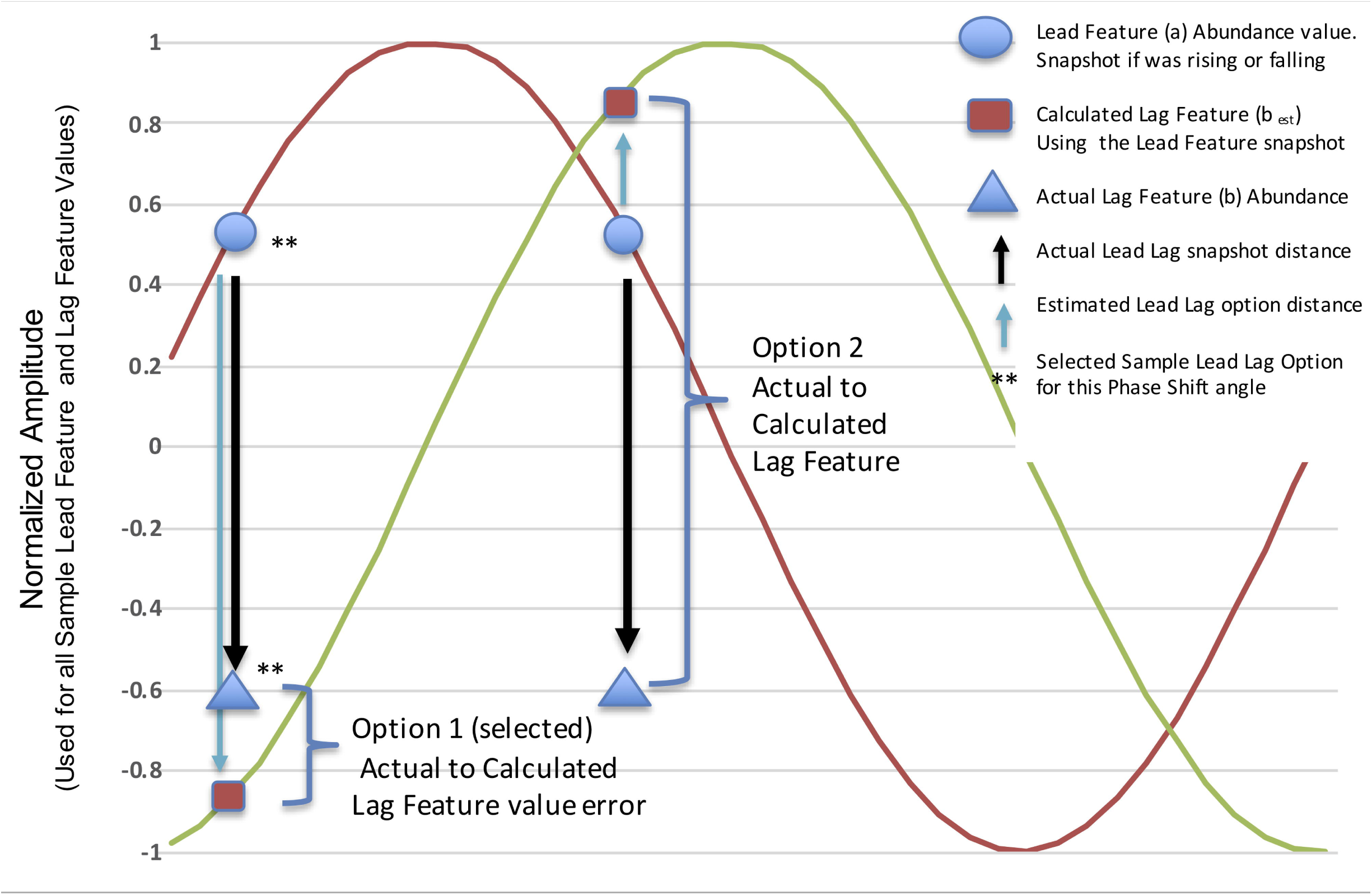
PSCrho determination. This figure only shows one step in the scenario for two features (a,b), from one sample, and for only one phase shift (90 degrees). The Lead feature (a) follows the red curve and the lag feature (b) follows the green curve. For every sample, each feature pair (a and b) will be tested, as both the lead and lag, for all Phase shift angles. The PSCrho is higher for the minimum overall total of actual lag feature minus estimated lag feature distance scores for this pair, where a leads b, or b leads a, across all phase shifts. The highest PSCrho score will identify the lead feature and the optimal phase shift angle.

The PSCrho has two components. The first is the correlation value (PSCrho) that is analogous to the Pearson rho. The second component is the phase shift (lag) in radians or degrees, which describes how the lag feature trails the lead feature.

The first step in applying the PSC algorithm is to normalize each feature’s values based on their data range across samples and set x_max_ = 1, x_min_ = -1. This will create a range for each feature of *x* ={-1:1}(Eq.1 and 2). To determine the PSCrho, the lead feature values are used as the acos(feature *x*_*i*_ value), and the phase shift is applied. Then, the cosine is applied to this phase angle to give the estimated value for feature *y*_*i*_ (Eq.3). The estimated feature *y* value and actual feature *y* values are used to calculate the correlation value (Eq.4).

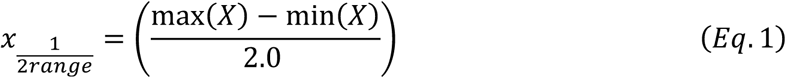

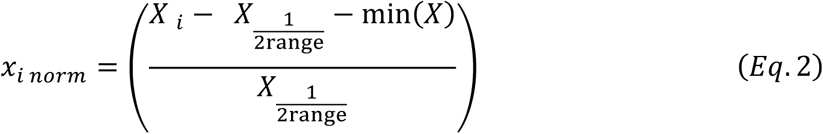

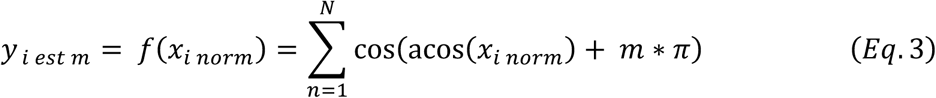

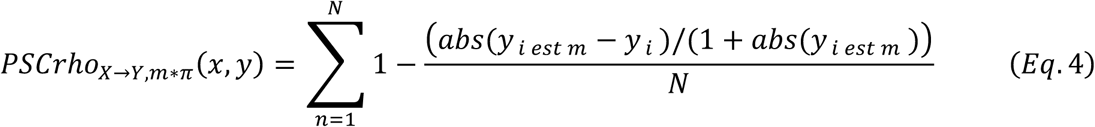

The PSC for a sample S_i_’s pair of feature actual values (*x*_*i*_,*y*_*i*_) determine the value *y*_*i*_’s expected value (y_i est_) based on the lead feature x_i_ being the correct value for the phase of the cosine wave being evaluated. The distance between *y*_*i*_ actual value and the *y*_*i est*_ is calculated. *y*_*i est*_ is the feature value if there were a perfect fit for both features with the pre-selected phase differential *m* \**π* (see following). The sum of the distance between each pair of *y*_*i*_ and *y* _*i est*_ is calculated and used in the PSC calculation in Eq.3

To calculate a PSCrho for each phase shift, run Eq. 4 to calculated between all *x*_*i*_ and *y*_*i*_ values for a given feature and the PSCrho for that phase shift is determined. Rerun this process for each phase shift increment. The phase shift increments go from *m* = 0 radians (no phase shift), to increments by 1/8 *π* radians to the final 1.0* *π* radians. To verify which feature would be the lead and the other the lag, all steps in Eq. 3&4 are repeated again, but with feature *y* leading and feature *x* trailing (swap *y*↔*x*).

From all the PSCrho values calculated, the largest *abs*(PSCrho) is the PSC that most represents the best possible relationship between the two features, whether feature *x* leads *y*, or *y* leads *x*, and by what phase lag increment. When the phase shift = -*π* (−180°) the rho becomes –rho for the inverse correlation determination. Feature x leading feature y by a non-zero, or non-180 degree value indicates possible causation of changes in x leads to changes in y.

The PSC alternative correlation approach effectively treats all sample feature collections as various points in time of the phase relationship between the two features in their domain.

For the situation with significant correlation where the phase shift is 0 degrees, the expected *normalized* values for feature *x* will roughly equal feature *y* for each sample. This is the same as a Pearson or Spearman correlation expectation. The wave function used in PSC is made to simulate a cosine wave by shifting all values by the difference of the max and min values for each Feature.

The PSC function uses the same scale as the linear version. A PSCrho = 1 indicates to features are perfectly aligned in a phase-difference sinusoidal relationship. A PSCrho = 0 does not necessarily indicate a sinusoidal relationship, while a PSCrho= -1 indicates a 180-degree inverse, possibly sinusoidal, relationship with the features moving inversely to each other. PSCrho is derived from performing both a zero phase (0-degree) shift relationship for each feature pair, and then a *π* phase (180-degree) correlation to test for an inverse relationship between the two features.

One PSC consideration is the removal of feature values that equal zero. The lead feature cannot influence the lag feature if the lead feature value is zero and the lag feature value is non-zero, since a value of zero implies that the lead feature is simply not present. The converse is true for zero lag feature values. Therefore, we remove all lead and lag pairs where one or both are zero. To calculate the PSCrho, both features are tested in the lead and lag roles without a priori knowledge of which feature will lead.. We removed sample feature values equal zero (no abundance) during the PSC Eq. 1 and Eq. 2 calculations. A zero value for the lead or lag feature indicates it was not present in its sample and didn’t participate in the correlation.

To ensure we have a balance of data values, we check for outliers, defined as a set of values that are less than 10% of all feature values being above or below the midpoint (0) will all the other feature values on the opposite side of the midpoint after data scaling Eqs 1 and 2. When the outlier limit is exceeded, we exclude that feature in any PSC determinations with other features.

To demonstrate the distinctions of phase shift correlations versus linear correlations, we ran the PSC algorithm with machine generated random values and determined the Pearson correlation for the same data. The comparison results for the PSCrho versus the Pearson rho are in Supplemental Table 1 and input data 2. PSC, similar to Pearson Correlation, cannot determine leading or lagging feature for both a 0 degree or 180 degree phase shift.

**Table 1.** Specific UC feature values that show the difference in the Pearson and PSC values. This table shows two specific feature pairs where the Pearson correlation is very low, but the PSCrho is high with a phase shift of greater than 22 degrees and less than 145 degrees. The full results are in S2

| Lead Feature | Lag Feature | Phase Shift Correlation | Lag Phase (degrees) | Pearson Correlation |
| --- | --- | --- | --- | --- |
| Lachnospiraceae_unknown_Blautia | Lachnospiraceae_Blautia | 0.873 | 22.5 | 0.844 |
| Ruminococcaceae_Ruminococcus | Lachnospiraceae_Lachnospiraceae_incertae_sedis | 0.865 | 45 | 0.701 |
| Lachnospiraceae_unknown_Robinsoniella | Ruminococcaceae_Faecalibacterium | 0.855 | 67.5 | 0.288 |
| Lachnospiraceae_Ruminococcus2 | Lachnospiraceae_Dorea | 0.851 | 45 | 0.656 |
| Erysipelotrichaceae_unknown_Coprobacillus | Lachnospiraceae_unknown_Roseburia | 0.841 | 45 | 0.725 |
| Lachnospiraceae_Roseburia | Lachnospiraceae_Fusicatenibacter | 0.839 | 45 | 0.626 |
| Lachnospiraceae_Roseburia | Lachnospiraceae_unknown_Lachnobacterium | 0.837 | 22.5 | 0.653 |
| Erysipelotrichaceae_Clostridium XVIII | Lachnospiraceae_Blautia | 0.837 | 67.5 | 0.311 |
| Erysipelotrichaceae_Clostridium XVIII | Lachnospiraceae_Lachnospiraceae_incertae_sedis | 0.836 | 22.5 | 0.73 |
| Erysipelotrichaceae_unknown_Coprobacillus | Lachnospiraceae_Blautia | 0.834 | 45 | 0.688 |
| Lachnospiraceae_Clostridium XIVb | Lachnospiraceae_Ruminococcus2 | 0.832 | 67.5 | 0.537 |
| Bacteroidaceae_unknown_Bacteroides | Ruminococcaceae_unknown_Clostridium IV | 0.832 | 135 | -0.817 |
| Ruminococcaceae_unknown_Clostridium IV | Bacteroidaceae_Bacteroides | 0.829 | 112.5 | -0.626 |
| Erysipelotrichaceae_Clostridium XVIII | Lachnospiraceae_unknown_Lachnobacterium | 0.827 | 45 | 0.799 |
| Erysipelotrichaceae_Clostridium XVIII | Ruminococcaceae_unknown_Faecalibacterium | 0.818 | 67.5 | 0.456 |
| Ruminococcaceae_unknown_Faecalibacterium | Erysipelotrichaceae_unknown_Coprobacillus | 0.817 | 45 | 0.657 |
| Lachnospiraceae_unknown_Anaerostipes | Lachnospiraceae_unknown_Roseburia | 0.813 | 22.5 | 0.529 |
| Lachnospiraceae_Clostridium XIVa | Lachnospiraceae_unknown_Clostridium XIVa | 0.813 | 22.5 | 0.596 |
| Lachnospiraceae_unknown_Lachnobacterium | Lachnospiraceae_unknown_Lachnospiraceae_incertae_sedis | 0.811 | 45 | 0.623 |
| Ruminococcaceae_unknown_Faecalibacterium | Ruminococcaceae_unknown_Flavonifractor | 0.81 | 67.5 | 0.207 |
| Streptococcaceae_Streptococcus | Lachnospiraceae_Blautia | 0.809 | 67.5 | -0.042 |
| Peptostreptococcaceae_unknown_Clostridium XI | Lachnospiraceae_unknown_Lachnospiraceae_incertae_sedis | 0.808 | 90 | 0.032 |
| Streptococcaceae_Streptococcus | Lachnospiraceae_unknown_Lachnospiraceae_incertae_sedis | 0.805 | 45 | 0.163 |
| Lachnospiraceae_unknown_Lachnobacterium | Lachnospiraceae_Lachnospiraceae_incertae_sedis | 0.805 | 45 | 0.525 |
| Lachnospiraceae_unknown_Anaerostipes | Lachnospiraceae_unknown_Lachnospiraceae_incertae_sedis | 0.802 | 22.5 | 0.606 |
| Lachnospiraceae_unknown_Roseburia | Lachnospiraceae_unknown_Dorea | 0.802 | 22.5 | 0.737 |
| Lachnospiraceae_Roseburia | Lachnospiraceae_Blautia | 0.802 | 67.5 | 0.094 |
| Streptococcaceae_Streptococcus | Lachnospiraceae_Lachnospiraceae_incertae_sedis | 0.799 | 45 | 0.296 |
| Lachnospiraceae_unknown_Anaerostipes | Lachnospiraceae_unknown_Ruminococcus2 | 0.796 | 22.5 | 0.738 |
| Lachnospiraceae_unknown_Ruminococcus2 | Lachnospiraceae_unknown_Lachnoanaerobaculum | 0.795 | 45 | 0.409 |

The goal of the PSC is to represent the relationship between biological features in a more natural way, where there are fewer values in the “between” state of two related features (metabolite quantities/microbe abundance) and more values in the full abundance or low abundance states. The PSC cannot determine the actual frequency for the phase shift relationship since we have no time periods to assign for the samples. The samples can be singular collections per participant and are assumed to be collected at different points in their microbiome cycle. Therefore, the Fourier transform cannot be applied to determine the frequency. However, the PSC applies a more natural constraint on the correlation by checking Feature A and Feature B as phase dependent values than as a linear relationship. There is likely a maximum frequency (minimum period) that this technique can address, for example, in intracellular features, where the rate of reaction may be accelerated by the proximity of reactants. This is unlikely to be the case in extracellular feature relationships. The rationale for applying Phase Shift Correlation to gut microbiome data is the possibility of causal relationships between microbial features of the microbiome. While causal relationships are difficult to prove due to complexity and numerous confounding factors in gut microbiome studies, presence of cyclically correlated features suggests possible mechanisms by which one species of bacterial community promotes or suppresses another one. For example, more metabolic pathways are altered in patients with IBD versus healthy control, than actual differences in microbial abundances (on genus level) between the two groups [8]. It is plausible to suggest that inflammatory pathways set in motion a series of events that lead to phasal differences in commensal gut bacteria.

## Results and discussion

We applied our PSC technique to the Ulcerative Colitis Study data [9] By applying the PSCrho along with the Pearson correlation (rho), we can see several additional relations via the PSCrho. A subset of data is presented in Table 1. The entire results are in S3 Table. We compared and visualized the results from the Pearson correlation and the PSC via Cytoscape (4). The nodes are measured UC sample features and the edges indicate correlation values as specified between the two features. A difference in Fig 3B is the PSC edges also have a phase relationship value (0 - 180 degrees or 0 - *π* radians) that indicates the phase shift between the lead and lag features (arrow points) towards the lag feature.

**Fig 3.**
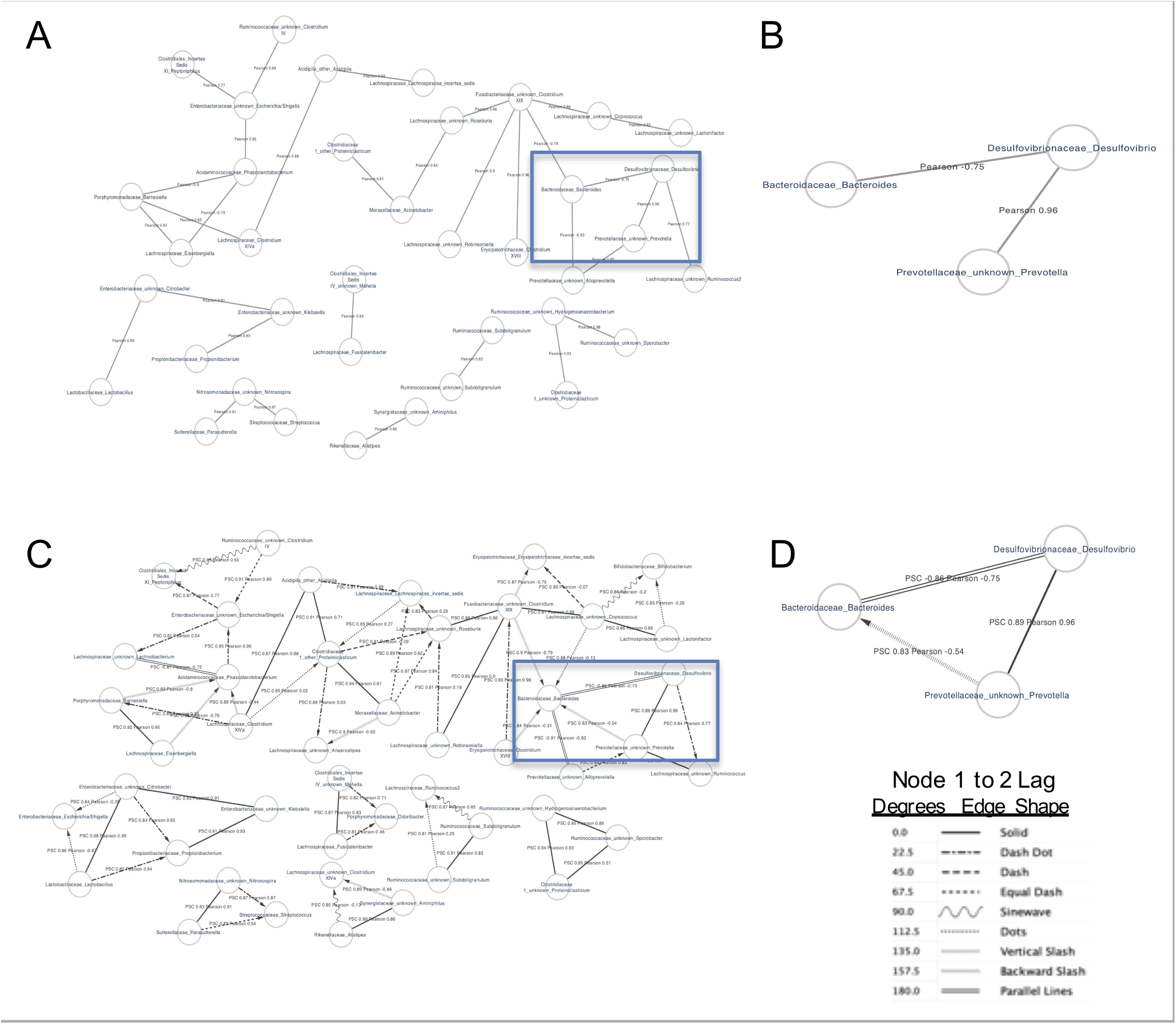
Pearson Correlation and Phase Shift Correlation (PSC) Network Diagrams. (A, B). UC cohort significant Pearson (A) and PSC (B) correlations in a subset of UC cohort data. The layout of nodes is preserved in both graphs. PSC shows additional edges not found by Pearson correlation. There is not a lead and lag feature for phase shifts of 0 or 180 degrees. The feature pair correlation value is listed on the edge connector. The location of the node, the link length and orientation are not significant. The outlined area focuses on a possible cyclic relationship between Prevotella, Desulfovibrio and Bacteroides. (C, D) ‘Correlation triangle paradox’ for Prevotella, Desulfovibrio and Bacteroides from the same UC cohort. Pearson correlation (C) was not significant between Prevotella and Bacteroides, however PSC (D) was significant and may show existing phasal relationship between the two genera.

If in both Fig 3A and Fig 3B, the layout of nodes representing bacterial taxa is preserved, it becomes apparent that the PSC captures relationships that are missed by Pearson correlations.

An example from the UC dataset is a possible cyclic relationship between *Prevotella, Bacteroides*, and *Desulfovibrio* (*Prevotella* – *Desulfovibrio*: 0° lag phase, PSCrho = 0.89, Pearson rho = 0.96, *Desulfovibrio* – *Bacteroides*: 180° lag phase, PSCrho = -0.86, Pearson rho = -0.75, *Prevotella* – *Bacteroides*: 135° lag phase, PSCrho = 0.83, Pearson rho = -0.54) Fig 3C. These genera represent two of the most significant phyla comprising intestinal microbiome: Bacteroidetes and Proteobacteria [10]. Healthy individuals and individuals with inflammatory bowel disease (IBD) have significant differences in relative abundances of Bacteroidetes, Firmicutes, and Proteobacteria, suggesting that cyclic correlation among them is possible [10]. Both *Prevotella* and *Bacteroides* are involved in mucosal glycoprotein degradation [11]. *Desulfovibrio*, sulfate-reducing bacteria, may aid glycoprotein degradation by mucin desulphation and co-occurs in samples with *Prevotella* [12]. This can be seen in our analysis with both PSCrho and Pearson correlations being significant (PSCrho = 0.89, Pearson = 0.96, and 0° lag). Moreover, a multi-study analysis of enteric bacterial profiles found that *Prevotella* and *Bacteroides* are frequently the only two taxa whose relative abundances exceed 40 %, and *Prevotella* tends to be inversely correlated with *Bacteroides*[13]. Our analysis found weak inverse Pearson correlation between the two genera (−0.54), but conversely a strong PSCrho (0.83) at 135° lag phase Fig 3D. Based on these findings, it is reasonable to say that our PSC triangle is well-supported.

## Conclusions

We present an alternative way to find correlations between two features based on the assumption of a sinusoidal abundance transition and a phase delay. PSC explains better than Pearson correlation some relationships, and helps explain the “dumbbell” effect, where the majority of data values are nearer to the min and max values that the middle. Supporting a sinusoidal relationship is the finding of high PSCrho values where the phase shift phase lag is in the 45 degrees < lag < 135 degrees range, where the linear correlation value is much lower. Additionally, the PSC addresses the correlation triangle paradox, where feature A is related to B, and feature B to feature C, but feature A is not related to feature C. PSC supports the possibility of assigning cause and effect to two features since, for non 0 or 180 degree phase shifts, the PSCrho assigns the lead and the lag feature to the relationship.

## Supporting information

1.S1.xlsx Raw UC Microbiome Data from Ref. 10

2.S2.xlsx All Pearson Correlation and PSCrho Correlation values for the S1 dataset.

4.S4.xlsx Examples of PSC concept based on random data generated in Excel

3.S3.txt Phase Shift Correlation python program source code to generate PSCrho values for the UC dataset in S1. PSC source code in Public Git Repo

## Supplemental Material

1. S1.xlsx Raw UC Microbiome Data from Ref. 10
2. S2.xlsx All Pearson Correlation and PSCrho Correlation values for the S1 dataset.
3. S3.py Phase Shift Correlation python program source code to generate PSCrho values for the UC dataset in S1. <u>PSC source code in Public Git Repo</u>
4. S4.xlsx Examples of PSC concept based on random data generated in Excel

